# Evolution of fast-growing piscivorous herring in the young Baltic Sea

**DOI:** 10.1101/2024.07.20.604447

**Authors:** Jake Goodall, Mats E. Pettersson, Ulf Bergström, Arianna Cocco, Bo Delling, Yvette Heimbrand, Magnus Karlsson, Josefine Larsson, Hannes Waldetoft, Andreas Wallberg, Lovisa Wennerström, Leif Andersson

## Abstract

The circumstances under which species diversify to genetically distinct lineages is a fundamental question in biology^1^. Atlantic herring is an extremely abundant zooplanktivorous species that is subdivided into multiple ecotypes that differ regarding spawning time and genetic adaption to local environmental conditions such as temperature, salinity, and light conditions^2,3^. Here we show using whole genome analysis that multiple populations of piscivorous (fish-eating) herring have evolved sympatrically after the colonization of the brackish Baltic Sea within the last 8,000 years postglaciation. The piscivorous ecotype grows faster, and is much larger and less abundant than the zooplanktivorous Baltic herring. Lesions of the gill rakers in the piscivorous ecotype indicated incomplete adaptation to a fish diet. This niche expansion of herring in the young Baltic Sea with its paucity of piscivorous species suggests that empty niche space is more important than geographic isolation for the evolution of biodiversity.

## Main

Genetic differentiation within a species and speciation requires some degree of reproductive isolation. This may occur due to geographic separation, where subpopulations accumulate genetic differentiation due to genetic drift and adaptation. The classical view of speciation was that it required the physical separation of populations^4^. However, more recent studies have documented that sympatric speciation may occur under certain circumstances^5^, for instance, as a result of hybridization between species, as shown for both plants^6^ and animals^7,8^.

Atlantic herring is an extremely abundant species with an estimated census population of 10^12^ (ref. 9). It is a zooplankton feeder with a key role in the North-Atlantic ecosystem as a link between plankton production and higher trophic levels (piscivorous fish, sea birds, and marine mammals). Atlantic herring is a broadcast spawner, where large schools of fish aggregate in coastal areas to spawn. Females and males release eggs and sperm into the water column before the fertilized eggs stick to vegetation or seafloor, where development occurs under local environmental conditions. Initial genetic studies in the 1980s suggested that all herring in the North Atlantic and the brackish Baltic Sea could be a single panmictic population because no genetic differentiation was noted for 13 polymorphic loci^10^. This was a surprising finding, given Linnaeus classified the Baltic herring as *Clupea harengus membras* in distinction of the Atlantic herring (*Clupea harengus harengus*) based on morphological differences^11^. The Baltic herring is smaller, with less fat than the Atlantic herring, most likely due to less productive plankton production and possibly physiological stress due to the low salinity in the Baltic Sea^12,13^, a young sea that has only existed for ∼8,000 years since the end of the last glaciation^14^. Recent studies based on whole genome sequencing confirmed a lack of genetic differentiation at selectively neutral markers but hundreds of loci underlying ecological adaptation related to variation in salinity, temperature, light conditions, and spawning time show genetic differentiation among ecotypes of Atlantic herring^2,3,15^. The *F_ST_* distribution, measuring the relative proportion of genetic variation within and between populations, deviates significantly from the one expected for selectively neutral alleles under a genetic drift model^16^, implying an important role for natural selection in determining allele frequencies.

This study aimed to explore genetic differentiation in a restricted area of the Baltic Sea (**Fig. 1a**), and in particular, to explore whether the herring denoted ‘Slåttersill’ by local communities constitutes a genetically unique population (**Fig. 1b**). Slåttersill is substantially larger than the dominating type of Baltic herring, where ‘sill’ refers to the Swedish name for the much larger Atlantic herring. This large type of herring has long been known to enter and spawn in coastal areas around hay harvest (Slåtter in Swedish) time in June. The presence of large herring in the Baltic Sea has also been noted by fishery biologists^17,18^, but it is unknown if such large herring just represent the tail of the size distribution of Baltic herring or a distinct subpopulation. Here, we show that Slåttersill is a genetically unique ecotype of herring with altered feeding behaviour and faster growth rate and that multiple subpopulations of such large, fish-eating herring exist in the Baltic Sea. This represents a remarkable example of sympatric differentiation and niche expansion in an extremely abundant broadcast spawner after colonizing a new environment within the last 8,000 years.

**Fig. 1.**
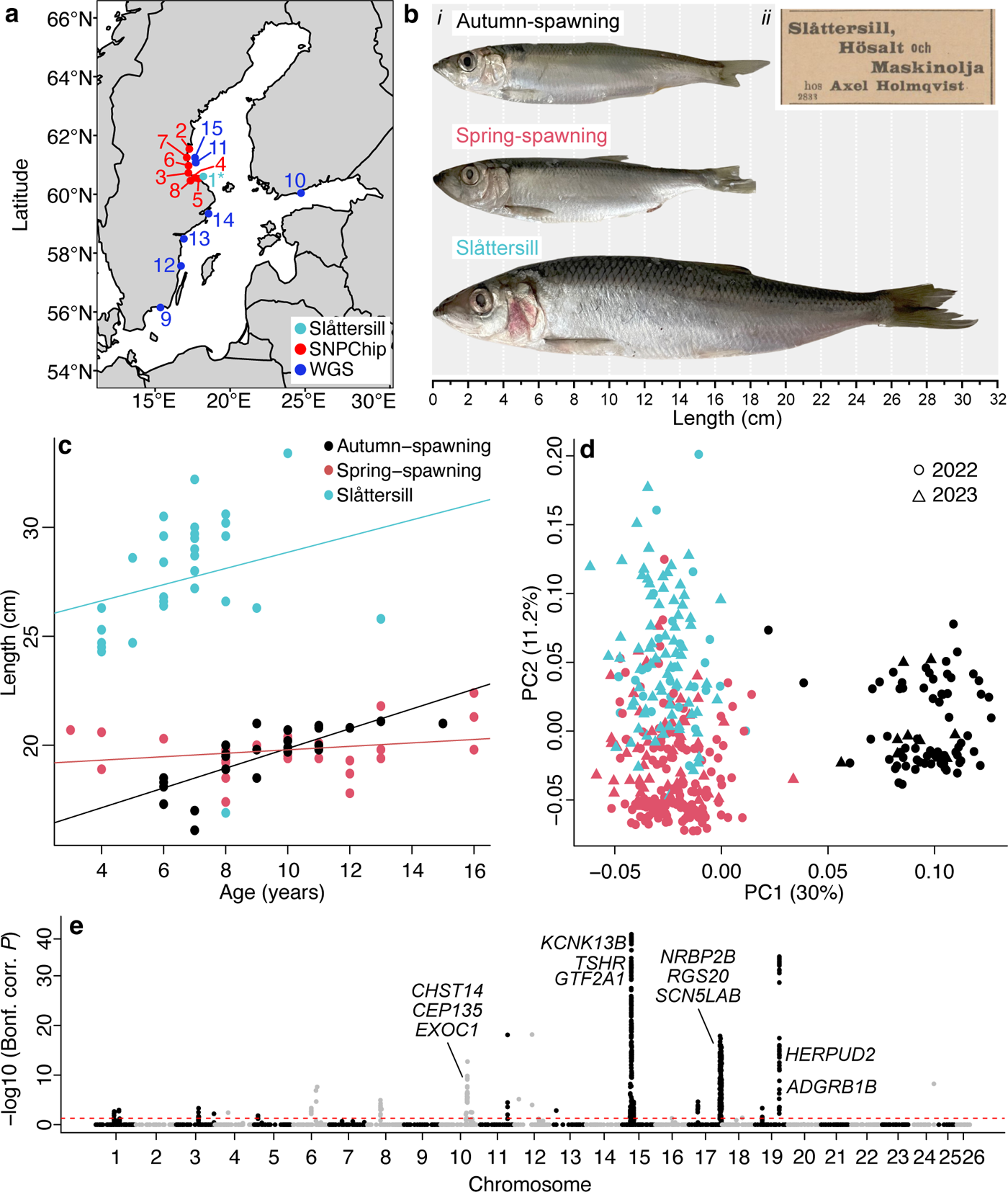
Characterization of the large Slåttersill and reference populations of small spring-spawning and autumn-spawning Baltic herring. (a) Map showing sampling locations in the Baltic Sea. Samples used for SNP-chip and whole genome sequencing is given in red and blue numbers, respectively. Sample 1 (light blue) is Slåttersill used for both type of analysis. Explanations for localities are in **Supplementary Table 1** and 8. (b) (i) Striking size differences between large Slåttersill and small spring-spawning and autumn-spawning Baltic herring. (ii) A newspaper clip showing an advertisement for ‘Slåttersill’ in the local newspaper Hudiksvallsposten from 1902-08-16. (c) Distribution of age and length among the three populations demonstrating differences in growth rate. (d) PCA plot based on genotype data for n=3840 SNP. Colours as in C. (e) Genome-wide, SNP-by-SNP contrast between Slåttersill (n=107) and small spring-spawning Baltic herring from the Bothnian Sea (n=201) utilizing 3,995 SNPs. x-axis: SNPs from individual chromosomes represented in alternating colours, y-axis: Bonferroni corrected *P*-values (-log10 transformed values).

### Slåttersill is a genetically distinct ecotype of Baltic herring

We collected multiple samples of both spring- and autumn-spawning planktivorous Baltic herring, as well as Slåttersill, at spawning from the same geographic region of the Baltic Sea (**Fig. 1a,b**, **Supplementary Table 1**). We used otoliths to determine the age of a subset of individuals from the three groups, which allowed us to establish growth curves (**Fig. 1c**). The data showed that the small spring- and autumn-spawning herring have similar growth curves, whereas Slåttersill has a faster growth rate, implying a different feeding behaviour. Slåttersill were also significantly younger at capture (2-3 years, t-test, *P*=0.001; **Fig. 1c, Supplementary Table 2**), demonstrating that Slåttersill are not simply older individuals at the tail-end of the small Baltic herring size distribution.

We genotyped all fish using a multi-species SNP-chip that includes ∼4,500 SNPs for Atlantic herring, including most regions of the genome showing genetic differentiation among ecotypes^19^. A principal component analysis (PCA) showed two major clusters representing genetic differentiation between spring- and autumn-spawning herring across PC1 (**Fig. 1d**). Slåttersill, which spawns in mid-June, cluster with the spring-spawners but with a clear shift across PC2 revealing genetic differentiation. Thus, Slåttersill is a genetically distinct population, and the two replicates of this ecotype from 2022 and 2023 fall very closely in the PCA plot.

We next made a genome-wide, SNP-by-SNP contrast between Slåttersill and spring-spawning Baltic herring (**Fig. 1e**). This revealed four regions with highly significant differentiation (χ^2^ test, *P*<10^-10^ after Bonferroni correction; **Extended Data Fig. 1**). The region on chromosome 17 overlaps one of four major inversions affecting ecological adaptation in Atlantic herring^3^. The regions on chromosome 15 and 19 include several loci previously associated with variation in spawning timing^3^. In addition to these four major loci, a handful of additional regions reached genome-wide significance (χ^2^ test, *P*<0.05). A list of all loci showing significant differentiation between Slåttersill and spring-spawning Baltic herring is in **Supplementary Table 3**.

### Is Slåttersill a fish-eating (piscivorous) ecotype?

The discovery that Slåttersill is a genetically distinct ecotype of Baltic herring prompted us to perform a detailed phenotypic comparison of Slåttersill, spring- and autumn-spawning herring, and Atlantic herring (**Supplementary Table 4**, **Extended Data Fig. 2**). We confirmed a previously reported difference^20^ that the number of vertebrae in the Baltic herring is smaller than that in the Atlantic herring (t-test, *P*<0.05), and the Slåttersill conformed with the Baltic herring by having the lowest average number (**Supplementary Table 4**). The number of gill rakers was higher in Baltic herring than in our samples of Atlantic herring, and also here, Slåttersill were more similar to Baltic herring than to Atlantic herring (t-test, *P*<0.05; **Supplementary Table 4**). The result is consistent with a previous study reporting higher gill raker counts in Baltic herring compared with most Atlantic herring populations^21^. Results from PCA of the morphometric data are given in **Supplementary Table 5** and **Extended Data Fig. 3**. PC I mainly reflects the overall size and explains 93% of the variation in the sample. PC II and the following components reflect variation less correlated to the size. Along the PC II axes, Slåttersill is most similar to the Baltic spring-spawning herring. Studies of individual measurement reveal that several characters, e.g., body depth, caudal peduncle depth, head length, cranial length, and eye diameter contribute to separation along the PC II axes. A larger head in Baltic herring is probably in part the result of fewer vertebrae giving a proportionally shorter body. Eye diameter is affected by growth rate^22^ and comparing specimens of similar size, Baltic herring (spring and autumn) have larger eyes than their Atlantic counterpart, possibly indicating the slower growth rate in the former^23^. For the large-sized Slåttersill, compared to Atlantic herrings of similar size, there is no distinct difference, possibly as a result of faster growth in Slåttersill (**Fig. 1c**), resulting in proportionally smaller eyes.

A striking difference was that 8 out of 10 Slåttersill showed gill rakers with clear lesions (**Fig. 2a-b**). The morphology of gill rakers in planktivorous (plankton-feeding) and piscivorous fish is drastically different because the former have numerous dense gill rakers for sieving plankton. Among clupeiforms, comparing *C. harengus* to the truly piscivorous Wolf-herrings (*Chirocentrus* spp.) serves as an example with the latter possessing a lower number, of short and sparsely distributed gill rakers^24^. Thus, a plausible interpretation of the lesions in Slåttersill is a recent switch from zooplankton to fish diet, not yet accomplished with genetic adaptation for piscivory.

**Fig. 2.**
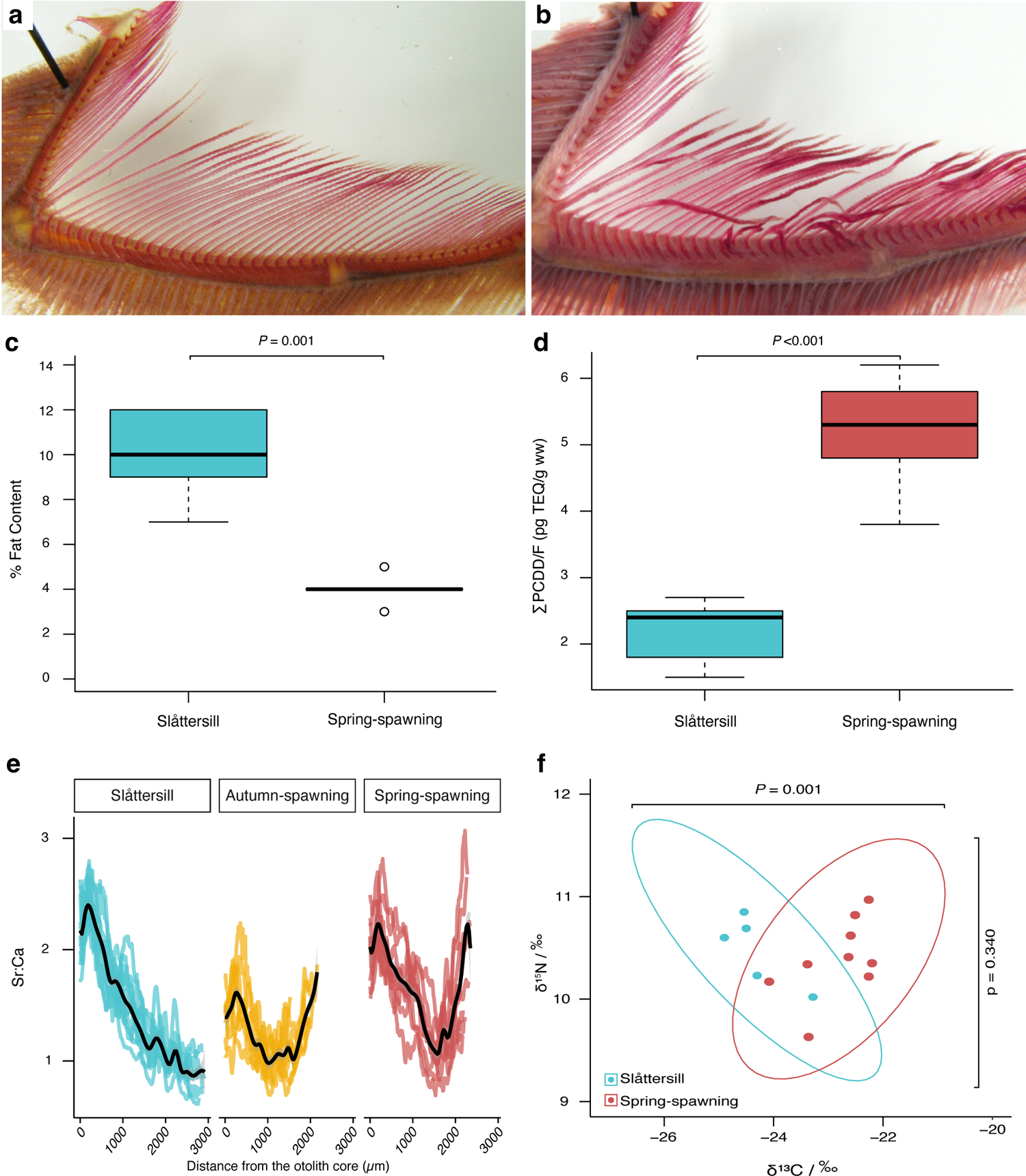
Phenotypic comparison of Slåttersill and small, spring-spawning Baltic herring. (**a**) Undamaged gill rakers from planktivorous spring-spawning Baltic herring. (**b**) Gill rakers with lesions frequently observed in Slåttersill most likely caused by a fish diet. (**c**) Fat content (%) in skeletal muscle from Slåttersill and small, spring-spawning Baltic herring. (**d**) Sum of polychlorinated dibenzo-p-dioxins (PCDD) and polychlorinated dibenzofurans (PCDF) measured in picograms of toxic equivalent per gram of wet weight per sample (∑PCDD/F (pg TEQ/g ww)), respectively, sampled from Slåttersill (n=98) and spring-spawning Baltic herring (n=170). (**e**) A comparison of lifelong otolith strontium:calcium profiles of Slåttersill (blue), autumn-spawning (yellow), and spring-spawning Baltic herring (red) from Gävlebukten with smoothed splines in black. (**f**) Stable isotope analysis of the same individuals as in **d** for carbon-13 to carbon-12 ratio relative to a standard (δ^13^C/‰), and for nitrogen-15 to nitrogen-14 ratio relative to a standard (δ^15^N/‰), expressed in parts per thousand.

Pollution from chlorinated organic compounds, e.g., polychlorinated dibenzo-p-dioxins and polychlorinated dibenzofurans (PCDD/Fs; dioxins), has been an environmental problem of concern in the Baltic Sea for decades, limiting the possibility for human consumption of herring and other fatty fish species^25^. Despite successful actions to reduce industrial effluent discharges^26^, the load of PCDD/Fs to the Baltic Sea is still significant, with atmospheric deposition of airborne pollutants from combustion processes being the dominating source^27^. Being fat-soluble, the dioxin content in fish generally correlates well with the lipid content when controlling for other factors, e.g., age. Surprisingly, Slåttersill, while having significantly higher lipid content than the spring-spawning herring (**Fig. 2c**; Wilcoxon rank-sum test, *P*=0.02), simultaneously had significantly lower PCDD/F levels in muscle and skin tissue (**Fig. 2d**, **Supplementary Table 6**; Wilcoxon rank-sum test, *P*=0.0005). As PCDD/Fs have a strong affinity to particles and accumulate in sediments, impacting benthic organisms in particular, a possible explanation is differences in the diet between ecotypes. Small planktivorous herring from the studied area, at least for parts of the year, feed on deep water macrocrustaceans, i.e., *Mysis,* containing relatively high levels of PCDD/Fs^28^. The much larger Slåttersill, as discussed above, most likely feeds on fish at larger sizes, likely being less exposed to the sediment-bottom water accumulation of PCDD/Fs. Moreover, the average age of Slåttersill was significantly lower compared to the small, spring-spawning herring (**Fig. 1c**, **Supplementary Table 2**), implying less time for PCDD/F to bioaccumulate, which also may have influenced the difference in PCDD/F levels^29^. However, levels of polychlorinated biphenyls (PCB), another bioaccumulating organohalogen of importance for the Baltic Sea, did not deviate between the two groups (**Supplementary Table 6**), making age differences likely less critical for the observed PCDD/F anomaly.

Next, we quantified the ratio of strontium (Sr) and calcium (Ca) in otoliths as a proxy for salinity^30^, to assess migration patterns. The results indicated lifelong ecotype-specific migration strategies (**Fig. 2e**). Pairwise comparison of otolith log transformed Sr:Ca slopes showed a general decline with age in adult Slåttersill, significantly different from the increasing trends in spring- and autumn spawning herring from Gävlebukten (**Fig. 2e, Extended Data Fig. 4,** and **Supplementary Table 7**). Adult Slåttersill otolith Sr:Ca profiles displayed both overall lower levels and less seasonal variations than spring- and autumn-spawning Baltic herring, suggesting that Slåttersill utilize habitats with lower salinity, and that Slåttersill do not make long-distance migrations (**Fig. 2e**). Thus, we conclude that the low levels of dioxin in Slåttersill are not caused by long-distance migration to feeding grounds with reduced contamination in the southern Baltic Sea, but is most likely explained by an altered diet and faster growth rate compared with small Baltic herring.

The altered growth rate, the damage on the gill rakers, and the significant reduction in dioxin contamination all implied an altered feeding behaviour and, most likely, a switch to a more nutrient-rich fish diet. To test this hypothesis, we performed a stable isotope analysis of the δ^15^N, the ratio of ^15^N/^14^N, which shows a positive correlation with the trophic level^31,32^. The comparison of Slåttersill and spring-spawning herring revealed no significant difference in δ^15^N (t-test, *P*>0.05; **Fig. 2f, Supplementary Table 6**). However, δ^15^N in the consumer tissue is also influenced by the protein content of its diet and by the growth rate of the consumer, both which can lower the so called trophic discrimination factor for ^15^N and result in lower δ^15^N values^33,34^. Thus, the fact that these two populations show a highly significant difference in growth rate (**Fig. 1c**) makes us suggest that Slåttersill, despite its similar δ^15^N ratio, is at a higher trophic level than the small spring-spawning herring, consistent with a switch to a high protein fish diet.

### Whole-genome sequencing reveals the presence of multiple subpopulations of piscivorous Baltic herring

The genetic constitution of the Slåttersill population was further characterized by whole genome sequencing, because the SNP-chip used here was designed prior to the discovery of this genetically unique population. We included six additional population samples of unusually large non-spawning Baltic herring from different parts of the Baltic Sea (**Fig. 1a**, **Supplementary Table 8**) to test if these represent the same population as Slåttersill. The stomach content of these large Baltic herring had previously been characterized (S. Donadi et al., submitted), and was found to be dominated by fish. Interestingly, the dominating prey (>60%) was three-spined stickleback (*Gasterosteus aculeatus*), which is highly abundant in the Baltic Sea and carries sharp spines that could explain the lesions noted on the gill rakers of Slåttersill (**Fig. 2b**).

Individual libraries for whole genome sequencing were prepared using Tn5 tagmentation^35^ and sequenced to an average 11.8X (±2.8) sequence coverage. A PCA analysis comparing the allele frequencies of seven population samples of these large piscivorous Baltic herring, including Slåttersill, with previously described ecotypes of Atlantic and Baltic herring^3^ revealed that the large Baltic herring clustered with spring-spawning Baltic herring or with some spring-spawning populations from the transition zone that connect the Atlantic Ocean with the Baltic Sea (**Fig. 3a**). Next, we made a genome-wide comparison of each large Baltic herring with the average allele frequencies observed in previously sequenced small, spring-spawning Baltic herring^3^. This analysis underscored differences between small and large herring and also demonstrated that the latter is genetically heterogeneous and comprises multiple subpopulations (**Fig. 3**; **Extended Data Fig. 5**). First, we identified a striking signal of genetic differentiation in a 3.9 Mb region on chromosome 20 (12.8–16.7 Mb) in five of the seven large herring populations (**Fig. 3b,c**; **Extended Data Fig. 5c-g**). This signal was present in Slåttersill but not detected in the SNP-chip analysis above since no SNPs from this region were present on the SNP-chip. The signal at chromosome 15 noted in the SNP-chip analysis of Slåttersill (**Fig. 1e**) was only replicated in the large herring from Gävlebukten, close to the area where Slåttersill was sampled (**Extended Data Fig. 5c**). The sharp signal on chromosome 19 (**Fig. 1e**), on the other hand was noted in all large herring except in the sample from Blekinge (**Extended Data Fig. 5b**). The most striking example of genetic heterogeneity among the large Baltic herring was the genetic differentiation for the 7.8 Mb supergene (inversion) present in three population samples, Stockholm, Östergötland, and Kalmar (**Fig. 3d**; **Extended Data Fig. 5e-g**).

**Fig. 3.**
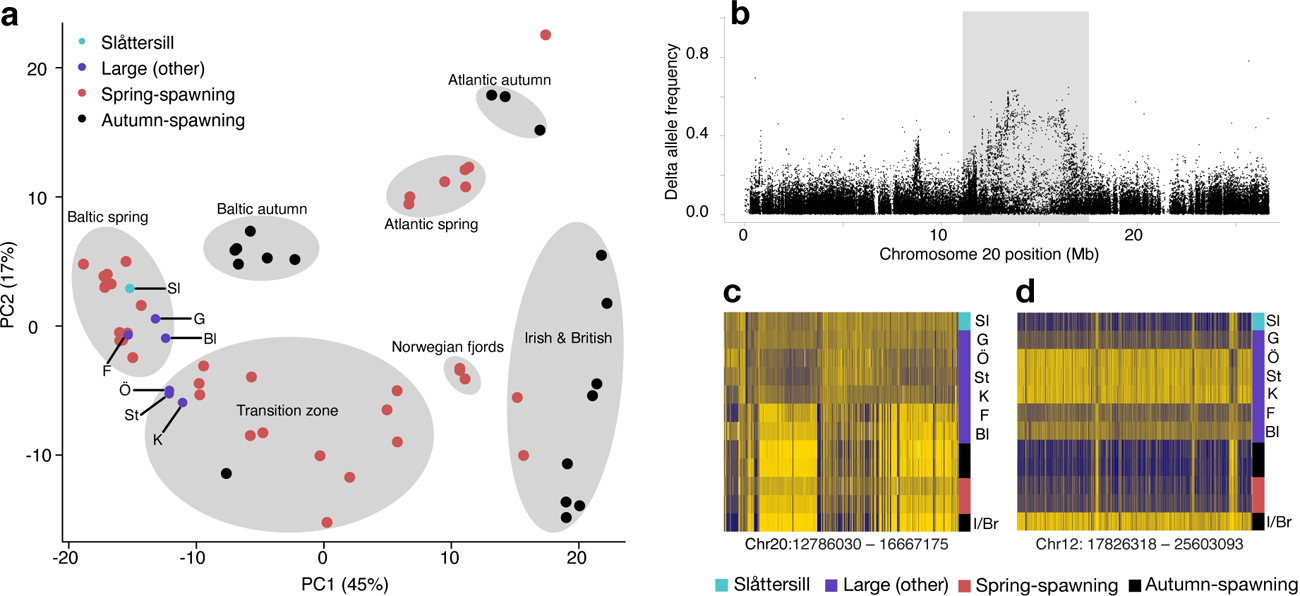
Whole genome sequencing reveals genetic heterogeneity among large Baltic herring. Comparison of Slåttersill and six other population samples of exceptionally large Baltic herring relative to previously sequenced Atlantic and Baltic herring samples^3^. (**a**) PCA plot of WGS data filtered to 407 (out of 794) markers previously used to define primary population groups Atlantic herring^3^. The seven primary population groups are represented as labelled grey ellipses, while the individual points represent pooled per-location averages. The colour of individual points represents the group each represents, i.e., Slåttersill (blue), large Baltic herring (other than Slåttersill, purple), spring-spawning herring (red), and autumn-spawning herring (black). (**b**) Delta allele frequency contrast of chromosome 20, between all large Baltic herring (including Slåttersill) and small, spring-spawning Baltic herring. (**c**) Zoom-in heatmap of the chr20 region of interest (20: 12.8 – 16.7 Mb) highlighted in the grey box in Fig. 3b, for all large Baltic herring, small spring- and autumn-spawning Baltic herring, and populations from Irish and British waters (I/Br) (delta allele frequency cutoff >0.25). (**d**) Zoom-in heatmap of the chromosome 12 inversion region (12: 17.8 – 25.6 Mb) for all large Baltic herring, small spring- and autumn-spawning Baltic herring, and populations from Irish and British waters (I/Br) (delta allele frequency cutoff >0.35). Population samples of large herring are labelled as follows: Sl = Slåttersill; G = Gävlebukten; Bl = Blekinge; F = Finland; Ö = Östergötland; St = Stockholm; K = Kalmar.

In conclusion, whole genome sequencing of seven population samples of large piscivorous herring reveals genetic differentiation compared with the much more abundant small planktivorous Baltic herring, but the piscivorous herring are composed of at least two distinct subpopulations. One is present in the Bothnian Sea, represented by Slåttersill and the sample from Gävlebukten (denoted northern piscivorous), and the other one was found in the central Baltic Sea and represented by the samples from Stockholm, Östergötland, and Kalmar (denoted southern piscivorous). Further work is required to determine if the population samples from Blekinge and Finland reflect additional genetically distinct populations of large piscivorous Baltic herring.

### Similar levels of nucleotide diversities among Baltic herring ecotypes but variable patterns of reproductive isolation

To further assess patterns of allele sharing, gene flow, and historical demographic trends, we grouped herring according to the four major ecotypes of the study region (northern vs. southern large piscivorous herring and autumn-vs. spring-spawning small planktivorous herring). We then used the whole genome sequencing data to estimate nucleotide diversity (Watterson’s *θ* and π per base), Tajima’s D, and long-term effective population sizes (*N*e) for each group (**Supplementary Table 9**). The data show very similar levels of diversity (*θ*_W_=in the range 0.45-0.50%). This is a somewhat surprising result given that it is obvious that planktivorous herring is much more abundant than the piscivorous herring in the Baltic Sea^18^. It implies that the establishment of piscivorous populations of Baltic herring was not associated with a strong founder effect, or that such a founder effect has subsequently been erased by gene flow. We performed pairwise estimates of genetic differentiation (*F*_ST_) among the ecotypes and estimated divergence to be in the range of 1.6–2.6%, with the large herring clustering with spring-spawners (**Extended Data Fig. 6a**). We found that the northern piscivorous herring (Slåttersill) shared much variation with spring-spawning planktivorous herring, as it showed lower genetic divergence to spring-spawners than any other ecotype across 47% of the genome and at 21 out of 26 chromosomes (**Extended Data Fig. 6b,c**). In contrast, the southern piscivorous herring appeared to share considerably less variation with spring-spawners, being most similar to the northern large herring (**Extended Data Fig. 6b– c**). These observations suggest that the northern and southern piscivorous populations have a common ancestry but the former may have exchanged more genetic material with planktivorous herring than the southern herring, which instead have been more reproductively isolated.

### A unique recombinant version of the Chr12 supergene in herring

One of the most important loci for local adaptation in herring is the 7.8 Mb supergene inversion on chromosome 12, for which haplotypes segregate according to latitude and temperature at spawning among populations^3,36^ Interestingly, this locus have diverged also among northern and southern piscivorous herring in the Baltic Sea. We found that the three piscivorous population samples (Stockholm, Östergötland, and Kalmar) representing the southern ecotype were almost fixed for a haplotype closely related to the Southern haplotype of the chromosome 12 supergene (**Fig. 3d**; **Extended Data Fig. 5e-g**). This haplotype is close to fixation in populations spawning in the waters surrounding Ireland and Great Britain, which is the warmest waters where herring spawning occurs. A comparison of southern piscivorous herring against spring-spawning planktivorous Baltic herring^3^, not carrying this haplotype, reveals the classical pattern for an inversion with strong genetic differentiation across the inverted region and with supersharp borders between SNPs inside and outside the inversion (**Fig. 4a**), whereas a comparison against herring from Ireland/Great Britain did not reproduce this inversion pattern (**Fig. 4b**), suggesting that the latter pair share very similar haplotypes. However, a closer inspection reveals three peaks of striking genetic differentiation within the inversion (marked **c**, **d**, and **e** in **Fig. 4b**). Region **c** apparently represents a recombination between the Northern and Southern haplotypes because the Southern haplotype present in large herring shares 12 diagnostic SNPs with Northern haplotypes (**Fig. 4c**); the differentiation in flanking SNPs most likely represents a selective sweep subsequent to the recombination event. Similarly, region **e** also represents a recombination between a Northern and Southern haplotype as indicated by many diagnostic SNPs (**Fig. 4e**), while region **d** may represent a selective sweep that has occurred on the Southern-like haplotype present in the southern piscivorous population (**Fig. 4d**). The presence of this unique version of the Southern haplotype, at a frequency of ∼0.8, which is rare or absent in the 60+ population samples previously sequenced^3^, provides evidence that the southern piscivorous population is a genetically unique population of herring not previously described.

**Fig. 4.**
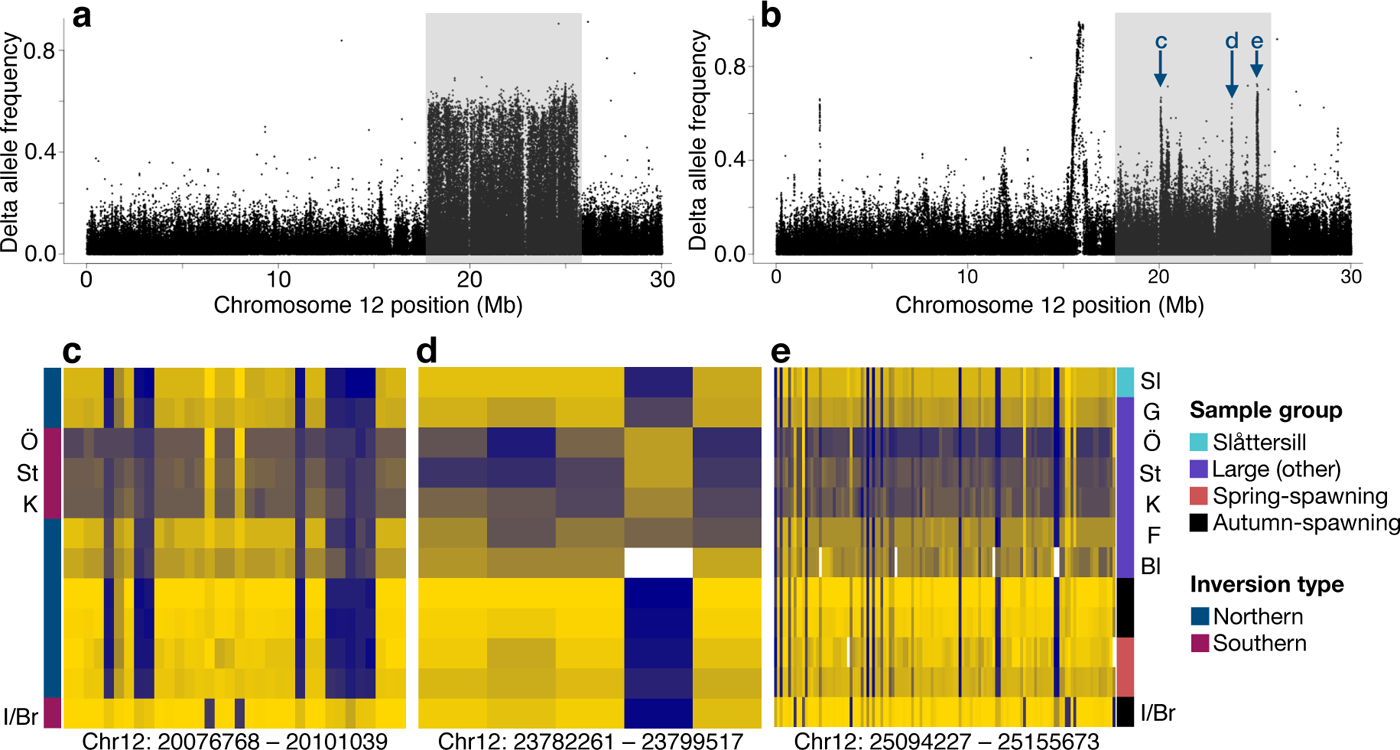
A recombinant version of the chromosome 12 supergene. (**a**) Delta allele frequency contrast between pooled allele frequencies for three large herring samples (Stockholm, Östergötland, and Kalmar) and small, spring-spawning Baltic herring. The chromosome 12 inversion region (12:17.8 – 25.6 Mb) is highlighted by the grey box. (**b**) Delta allele frequency contrast between pooled allele frequencies for three large herring samples (Stockholm, Östergötland, and Kalmar) and Atlantic herring from Irish and British waters. Significant peaks of differentiation within the chr12 inversion region (grey box) are annotated relative to the subsequent zoom-in heatmaps used to visualize each peak. (**c-e**) Zoom-in heatmaps of the chr12: 20,076,768 – 20,101,039 region (**c**), the chr12: 23,782,261 – 23,799,517 region (**d**), and chr12: 25,094,227 – 25,155,673 region (**e**). The colour code to the left of (**c**) indicates if the predominant inversion haplotype is classified as southern or northern^3^. The population samples are coloured as follows: Slåttersill (blue), other large Baltic herring (purple), spring-spawning herring (red), and autumn-spawning herring (black). The labelling of large Baltic herring samples is as in Fig. 3, where: Sl = Slåttersill; G = Gävlebukten; Ö = Östergötland; St = Stockholm; K = Kalmar, F = Finland; Bl = Blekinge; I/Br = Ireland and Britian.

## Discussion

This study has revealed an unexpected sympatric genetic differentiation and niche expansion in the Baltic herring. The Atlantic herring, as well as the small Baltic herring, are typical plankton-feeding fish of major importance to the ecosystem, both displaying gill raker morphology consistent with this feeding behaviour (**Fig. 2a**). Atlantic herring in proper marine environments may occasionally feed on fish eggs and larvae^37,38^, but extensive feeding on larger fish has not, to the best of our knowledge, been described outside the Baltic Sea. The common small Baltic herring feed on zooplankton (90%), mysids, and benthos^39^, while it has been reported that unusually large Baltic herring may feed on fish^17,18^. However, it was unknown if these herring simply represented the right-most end of the size distribution, and that Baltic herring of a specific size may switch feeding behaviour. This study demonstrates that these herring constitute genetically differentiated ecotypes with a distinct growth and migratory pattern (**Fig. 1, 2e and Extended Data Fig. 4**).

The finding that herring shows niche expansion in the Baltic Sea, but most likely not in the Atlantic Ocean, likely reflects reduced competition from other piscivorous fish in the Baltic Sea. The Baltic Sea is a young water body formed within the last 8,000 years, after the end of the last glaciation^14^. Few marine fish have colonized the Baltic Sea and large predators such as different species of mackerel, tuna, and pelagic gadoids are not present. Several other species of fish show adaptation to the Baltic Sea, such as Atlantic cod^40^, which has differentiated strongly from its conspecifics in fully marine environments, and the endemic Baltic flounder^41^. However, the ecotypes of herring described here appear to be unique by showing both sympatric genetic differentiation and niche expansion.

Sympatric differentiation and niche expansion are well documented in freshwater fish, in particular salmonids^42^. For instance, in the Arctic char it is described that morph separation may involve three different phases starting with differentiation due to plasticity but no reproductive isolation, followed by genetic differentiation and incomplete reproductive isolation, and eventually complete reproductive isolation^42^. Similarly to both salmonids and the radiation of African cichlids^43^, the Atlantic herring has diverged into multiple ecotypes exploiting different environments and food resources.

We did not notice significant genetic differentiation at neutral loci in the piscivorous Baltic herring despite the striking genetic differentiation at dozens of adaptive loci (**Fig. 1e, 3 and 4**), suggesting low genetic drift and incomplete reproductive isolation, although to a different degree between the northern and southern piscivorous populations. Population genetics theory has established that very little gene flow (one migrant per generation) is sufficient to avoid differentiation and loss of heterozygosity in subdivided populations^44^. The level of gene flow that has eliminated differentiation at neutral loci has not prevented the sympatric evolution of piscivorous Baltic herring, implying that this ecotype has evolved through strong natural selection. What makes the evolution of piscivorous Baltic herring unique is that this niche expansion has not been reported in other parts of the species distribution and the conundrum why the piscivorous ecotypes have not been genetically absorbed by the abundant planktivorous Baltic herring with an estimated census population size of ∼70 billion individuals^9^.

How is sympatric genetic differentiation possible in an extremely abundant fish that makes long-distance migrations, and is a broadcast spawner? Our recent description^45^ of a long-standing hybrid population formed by hybridizations between the two sister species, Atlantic herring and Pacific herring (*Clupea pallassii*), implies that there is not any strong prezygotic isolation in herring, as expected for a broadcast spawner lacking mate choice. The fact that this Pacific/Atlantic hybrid population has existed for thousands of years without being absorbed by the huge population of Atlantic herring present in the same geographic area is of interest in relation to the ability of the piscivorous populations of Baltic herring to maintain their unique genetic profile. This occurs even though they are spawning in the same geographic region during the same period as the spring-spawning planktivorous herring. The implication is either that the herring has a more sophisticated homing behaviour than previously thought^46^, or, alternatively, there exists a spawning behaviour that restricts gene flow from other populations. A strong homing behaviour is adaptive in herring because it allows genetic adaptation to environmental conditions at spawning. The herring deposits the fertilized eggs on vegetation or on the sea floor, and these are thereby exposed to the environmental conditions on the spawning grounds to a much larger extent than a pelagic spawner where the fertilized eggs drift with sea or ocean currents. Further, there are very strong environmental gradients (e.g. temperature, salinity, water transparency, and primary production) in the Baltic Sea coastal zone, especially during spring^47^ which would underscore the importance of a pronounced homing behaviour for populations to optimize chances of successful reproduction.

The main prey of the piscivorous Baltic herring appears to be the abundant three-spined stickleback, a species with sharp spines known to cause damage to its predators^48^. Similar lesions have been documented in fish-eating kokanee salmon (land-locked *Oncorhynchus nerka*) that also had stickleback as its main prey. While morphological, genetic, and stable isotope techniques did not support the separation of piscivorous and planktivorous kokanee into discrete ecotypes, it was suggested that an increase in gill raker spacing due to a larger body size of piscivorous individuals, in combination with damage on gill rakers, would reduce foraging efficiency on zooplankton and thus reinforce the shift to piscivory on the individual level^49^. A similar phenotypic plasticity and feedback mechanism may have been the initial step for the evolution of the piscivorous herring ecotype, followed by a certain degree of reproductive isolation due to strong homing behaviour and natural selection.

The discovery that a genetically distinct subpopulation of large, fish-eating Baltic herring is close to fixation for a unique recombinant version of the Southern haplotype for the chromosome 12 inversion demonstrates that this supergene undergoes dynamic evolution. This is consistent with a detailed characterization of the four major supergenes in herring (on chromosome 6, 12, 17, and 23) based on PacBio long-read sequencing that reveals that these originated more than a million years ago and that there is a considerable gene flux between inversion haplotypes at all four loci^50^. These supergenes play a prominent role in ecological adaptation in Atlantic and Baltic herring, and the supergenes on chromosome 17 (**Fig. 1e**) and chromosome 12 (**Fig. 4**) have both contributed significantly to the genetic differentiation of the piscivorous Baltic herring. The Southern haplotype is uncommon at higher latitudes in both the Baltic Sea and the Atlantic Ocean^3,36^, hinting at parallel sorting of adaptive alleles at this locus between ocean basins.

The current rapid loss of biodiversity and genetic diversity due to human expansion and exploitation of natural resources are of major concern^51^. This study demonstrates that also in extremely abundant species, precious and potentially vulnerable genetic diversity may be present, although not detectable using the standard approach for monitoring genetic diversity using a sparse set of selectively neutral markers. There is an obvious risk that the currently high fishing pressure on Baltic herring populations may lead to losses of genetic diversity and unique subpopulations, warranting cautious management. This could be particularly important for the Baltic Sea, where the ecosystem depends on few species and trophic interactions are easily disturbed. For example, perch and pike normally prey on coastal three-spined stickleback but have recently declined, leading to massive increase in sticklebacks that instead feed on the larvae of their predators (so-called predator–prey reversal), limiting their recruitment and reinforcing trophic regime shifts in the Baltic Sea^52^. If managed favourably, the large piscivorous herring characterized here could help control the stickleback population and be an unexpected and valuable ally in restoring Baltic food-webs. A comprehensive understanding of the population structure is particularly important in species that are heavily exploited by industrial fishing^19^, both for avoiding unsustainable overfishing and for protecting ecosystem function. This includes the Atlantic and Baltic herring, which sustain one of the top ten most important fisheries in the world^53^.

## Methods

### SNP-chip analysis

Local fishers were chartered for capture of fish from eight locations across Gävlebukten, Sweden (**Supplementary Table 1**). Charters were conducted in 2022 and 2023, between April and October to capture both Slåttersill (n=107), as well as spring-(n=201) and autumn-spawning Baltic herring (n=79). Length and weight were recorded before collecting muscle tissue using a Biomark Tissue Sampling Unit (Biomark LLC), then stored at -80°C.

DNA extraction and the following SNP-chip analysis was performed by IdentiGEN (Dublin, Ireland) using the MultiFishSNPChip_1.0 array (FSHSTK1D)^19^. The herring component of the SNP-array was designed to cover all independent regions of divergence, that had been identified at the time, including a large proportion of missense mutations showing strong genetic differentiation between populations. The array also includes a set of neutral markers, defined as having minor allele frequency above 0.3 but little inter-population variation spread approximately evenly along the chromosomes. The full list of SNP-designs is provided as **Supplementary Table 10**.

The genetic spawning season for all fish was predicted using AssignPop^54^. All SNP-chip genotyped fish were compared against an Atlantic herring reference database comprising n=8 individuals simulated from per-location (n=53) pooled allele frequency data^3^. Each of the 53 reference pools were used previously to define the seven population clusters inherent to the species’ North Atlantic range (i.e., Baltic Sea (Autumn), Baltic Sea (Spring), Atlantic Ocean (Autumn), Atlantic Ocean (Spring), Ireland and Britain, Norwegian Fjords, and Transition Zone)^3^, with the current Atlantic herring reference database reflecting these assignments. All SNP-chip genotyped fish were therefore assigned to one of the seven population clusters in AssignPop, with genetic spawning season characterized only when individuals were assigned to either Baltic Sea (Autumn), Baltic Sea (Spring), Atlantic Ocean (Autumn), or Atlantic Ocean (Spring) groups. Population assignment was deemed successful when AssignPop assignment probabilities were >66% for a single population cluster (i.e., two-thirds more likely than the next alternative outcome^55^).

Principal component analysis was carried out using function (--pca) in PLINK2^56^. The MultiFishSNPChip_1.0 array contains a high redundancy of SNPs within inversions, showing strong linkage disequilibrium. Therefore, PCAs were both calculated, including all available SNPs, or a dataset in which the inversions were treated as single loci. To achieve the latter, all individuals were genotyped for the inversion, by comparing SNP data against reference haplotype assignments sourced from Han *et al.*^3^. Reference haplotypes constituted the consensus allele for either the northern (*N*) or southern (*S*) haplotype derived from pure northern (Norway Spring, Iceland Spring, and Greenland Summer) or pure southern haplotype (Isle of Man Autumn, Celtic Sea Autumn, and Downs Winter), respectively. Each SNP within inversion was scored against the reference set, and genotyped as *NN*, *NS*, or *SS*. The genotype for each inversion was then deduced (*NN, NS*, or *SS*) consistent with the predominant haplotype across each inversion. As the spawning season is expected to be the predominant signal differentiating populations of Baltic herring, individuals were visualized in R, and assignments to spawn season were predicted using K-means clustering (K=2) of PLINK-derived eigenvalues.

Allele frequencies were calculated using PLINK2^56^ for all population samples independently, as well as for groups of spring-spawning Baltic herring, autumn-spawning Baltic herring, and Slåttersill fish groups. PCA was calculated in R (*prcomp* function) and visualized using ggplot2^57^. Genotype BED files and association analyses were generated with PLINK (v1.9), and Bonferroni corrected P-values calculated using the --assoc and –adjust functions, then visualized using R. Gene annotation for all SNP was sourced from BioMart^58^ using the ’charengus_gene_ensembl’ dataset as a reference.

### Whole genome sequencing (WGS)

The population samples of exceptionally large Baltic herring included, in addition to Slåttersill, five population samples from Sweden and one from Finland (**Supplementary Table 8**). A control sample (Gävleborg Norrsundet) was also collected, representing small, spring-spawning Baltic herring from the same geographic region as where Slåttersill occurs. All fish were sampled between 2020 and 2022, with fish length recorded before muscle tissue was collected by dissection and stored at -80°C.

A total of 86 individuals was used to prepare WGS libraries with Tn5-based tagmentation. The method for library preparation was an implemented version of a previous protocol used in our laboratory^59^ adapted from Picelli *et al*.^35^. DNA was extracted with the DNeasy 96 Blood & Tissue Kit (Qiagen, Hilden, Germany) and quantified with the NanoDrop One C (Thermo Fisher Scientific, Madison, Wisconsin, USA). Genomic DNA from Slåttersill was extracted at IdentiGEN Ltd. (Ireland) as part of the SNP-chip analysis and consisted of a crude extract obtained using Chelex. The DNA integrity for all samples was checked on a gel. The samples were then diluted to 10 ng/µL and 2 or 4 µL were used for library preparation. Tn5 tranposase (Tn5 Tnp) was purchased from the Protein Science Facility at the Karolinska Institute (Stockholm, Sweden). The enzyme was loaded with pre-annealed mosaic end primer pairs, ME-A (Fw: 5’-TCGTCGGCAGCGTCAGATGTGTATAAGAGACAG-3’and R: 5’-CTGTCTCTTATACACATCT-3’) and ME-B (Fw: 5’-GTCTCGTGGGCTCGGAGATGTGTATAAGAGACAG-3’and R: 5’-CTGTCTCTTATACACATCT-3’), for 2h at room temperature. The complex was diluted to a final concentration of 8 or 6.4 ng/µL in the tagmentation reactions. Tagmentation was allowed to proceed for 5 to 10 min at +55°C, then Tn5 transposon was stripped off the DNA template with 0.04% SDS. Primer sequences for library indexing were from Illumina Nextera Library Prep Kits and were combined to give a unique combination of i5 and i7 indexes per sample. The primers were appended to the DNA molecules by a 9-cycle PCR with the KAPA HiFi PCR Kit (Kapa Biosystems Pty, Cape Town, South Africa) with cycling parameters of 72°C for 3 min for gap-filling and 98°C for 30 s for denaturation, followed by 9 cycles of 98°C for 30 s, 63°C for 30 s, 72°C for 3 min. A double-sided size selection of 0.5X-0.7X was performed with AMPure XP paramagnetic beads (Beckman Coulter, Inc., Brea, California, USA)^60^. An aliquot of each individual library was mixed with 70 µL of Qubit HS reagents (Life Technologies Corporation, Eugene, Oregon, USA) prepared according to instructions and fluorescence was read with a TECAN infinite M200 microplate reader (Tecan Austria GmbH, Grödig, Austria). The Qubit controls were included in the measurements to construct a linear regression line and calculate the individual library concentrations. Three ng were pooled per each library and the pool was concentrated to 35 µL with another round of 0.5X-0.7X size selection. The insert size was checked with a TapeStation 4150 (Agilent Technologies, Waldbronn, Germany), and the concentration measured by qPCR with the KAPA Library Quant Kit (Kapa Biosystems Pty, Cape Town, South Africa). Sequencing consisted of paired-end sequencing with 150 bp read length and was performed by the SNP&SEQ Technology Platform in Uppsala on a NovaSeq 6000 with a S4 flowcell and v1.5 sequencing chemistry.

Raw paired-end reads were mapped to the *Clupea harengus* reference assembly ^36^(Ch_v2.0.2.fasta) using samtools^61^. Variants were called from BAM files using a standard gatk pipeline^62^. Firstly, BAMs were indexed using samtools, before individual gvcf were generated using the *gatk HaplotypeCaller* function. All sample-specifc gvcf were combined using *CombineGVCFs* and genotypes called using *GenotypeGVCFs*. SNP and INDELs were subset into independent variant files using *SelectVariants*, with the SNP-specific subset further filtered for biallelic SNP and multiple expression using *VariantFiltration* (QD <2; QUAL < 30; SOR >3; FS > 60; MQ <40; MQRankSum < -12.5; ReadPosRankSum < -8) and bcftools/1.17. Pooled allele frequencies were calculated independently using PLINK2^56^ for all populations.

Allele frequencies per sample group were calculated from the called genotypes using the “freq2” output from vcftools, v0.1.16^63^, and delta allele frequencies were then calculated using R v3.6^64^. For plotting purposes, only sites with at least 140 out of a possible 152 (92%), observed chromosomes were retained. SNPs were imputed and phased with BEAGLE v4^65^ ahead of estimation of levels of genetic diversity and divergence among ecotypes. Samples were then grouped into ecotypes according to location, morphology and spawning genotypes (**Supplementary Table 9**). Levels and patterns of diversity (*θ*_W_, π and Tajima’s D) were estimated across the 26 chromosomes using a custom Perl script that incorporated BioPerl code for Tajima’s D^66,67^. The standard equation *N*_e_=*θ*_W_/4μ was used to infer effective population size using per-base *θ*_W_ estimates and the herring mutation rate of 2.0×10^-9^ per base and generation^9^. Reynold’s *F* ^68^ was used to estimate pairwise divergences among ecotypes across the whole genome and in 10 kb windows and the UPGMA clustering algorithm was used to build a population tree from genome-wide estimates, as implemented in the neighbor program in PHYLIP v3.697^69^.

### Otolith analysis

The individual ages of samples of Slåttersill (n=32), spring-spawning Baltic herring (n=31), and autumn-spawning Baltic herring (n=26) were estimated by counting annuli on transverse otolith sections stained with toluidine blue.

For otolith chemistry analyses, otoliths of Slåttersill (n=10), spring-spawning Baltic herring (n=10), and autumn-spawning Baltic herring (n=8) were first embedded in epoxy, then hand-grinded and polished to the sagittal midplane to expose the core. Otolith trace elemental concentrations of strontium (^88^Sr) and calcium (^43^Ca) were analysed with laser ablation inductively coupled plasma mass spectrometry (LA-ICP-MS) at the College of Environmental Science and Forestry at the State University of New York (SUNY-ESF) in Syracuse, New York, USA. Otolith material was ablated along a line transect from the core (birth) to the anterior rostrum edge (death) following the maximum growth axis (**Extended Data Fig. 4a**). The otoliths were analysed with a round laser spot size of 85 µm at a scanning speed of 5-6 µm/sec with a repetition rate of 10 Hz, using helium and argon as carrier gas. The reference material MAPS-5 was used as the primary standard and NIST SRM 612 for tuning and checking the daily performance of the instrument. Iolite, Excel, and R were used for data reduction and statistical analyses with α < 0.05 as the threshold for statistical significance^70–72^.

Repeated measurements of Sr:Ca along the line transect of each otolith were used for performing mixed-effects regression analyses (lmer) with the R package lme4^73^. Otolith Sr:Ca was not normally distributed and therefore log transformed logarithmically. Spawning type, distance from the otolith core (µm) and the interaction term were used as fixed factors and sample ID as the random factor (model formula: log Sr:Ca ∼ spawning type + distance + (spawning type * distance) + (1| sample ID)). No multicollinearity between predictor variables was detected (variance inflation factor (VIF) < 5). Statistical pairwise comparisons of migration patterns of adult Slåttersill, spring-spawning Baltic herring, and autumn-spawning Baltic herring were performed by comparing the model slopes of log (Sr:Ca) at ≥1500 µm from the otolith core corresponding to adult fish using the emtrends function in the emmeans R package^74^. The R package ggplot2 was used for graphical comparisons of spawning types^57^.

### Morphological analysis

A total of 47 specimens were analysed including 10 specimens each of Slåttersill and Baltic spring-spawning herring (**Supplementary Table 4**). The material was received frozen to the Swedish Museum of Natural History (NRM), subsequently thawed, fixed in formalin, and transferred to 70% ethanol before analyses. Measurements and counts were obtained from x-rays or with digital calipers and rounded to the nearest 0.1 mm (**Extended Data Fig. 2**). Counts of vertebrae include the Atlas vertebra and the last half centrum (urostyle) being part of the caudal fin skeleton. This method yields higher counts, plus one vertebra, compared to conventional methods. Counts of dorsal and anal fin rays were estimated by counting their supporting pterygiophores from X-rays. The right side outermost gill arch was carefully dissected from each specimen and dyed with Alizarin red^75^ to obtain total gill raker counts. The length of the second uppermost gill raker on the lower limb was measured with a reticle eyepiece mounted on a stereomicroscope. Principal component analyses (PCA) on log-transformed measurements were used as an ordination method^76^. PCA was performed using SYSTAT 13. Gill raker length was excluded from PCA due to missing data for several specimens of Slåttersill due to the observed damages (**Fig. 2b**) and the lack of any obvious difference between samples.

### Organochlorine content and stable isotope analysis

After removing the head and guts, approximately ten individuals were homogenized to a mixed sample containing muscle tissue, bones, fins, skin, and subcutaneous fat, following the European Union guidelines (EU 2017/644) for food safety analyses. Five and ten mixed samples were prepared for Slåttersill and small spring-spawning herring, respectively. 100 g from each mixed sample was transferred to a glass jar, frozen, and transported to ALS Global laboratory in Prague, Czech Republic. After Soxhlet extraction, lipid content and levels of PCDD/Fs and PCB were quantified using high-resolution gas chromatography (HRGC) following US EPA 1613 (PCDD/F) and US EPA 1668 (PCB).

After freeze drying of the homogenized mixed samples, stable isotope analysis was performed with an Elemental Analyzer - Isotope Ratio Mass Spectrometer (EA-IRMS) by the SLU Stable Isotope Laboratory (Swedish University of Agricultural Sciences) with standard procedures and using the same samples as used for measuring organochlorine content.

## Data availability statement

Sequence data in this study have been deposited in the Sequence Read Archive under BioProject PRJNA642736.

## Code availability statement

All custom code is available in https://github.com/LeifAnderssonLab/Piscivorous_Baltic_Herring.

## Acknowledgments

We thank everyone who contributed to the collection and sampling of herring, Lars-Ivan Hållstrand (Hästskär) for collection of fish and for informing us about the existence of the Slåttersill population, Christin Appelqvist, Kerstin Johannesson, and the crew at SD511 EROS III for Atlantic herring caught as dead by-catch that we could use for morphological examination, Florian Berg for samples of Atlantic herring from Norway, and Identigen Ltd. for providing extracted genomic DNA. Sincere thanks are also due to Karin Limburg, Debra Driscoll and Marju Kaljuste for help conducting laboratory analyses, Agnes Karlsson (Stockholm University) for advice on stable isotope analyses and data interpretation, to Anton Larsson for finding the Slåttersill advertisement, to Peter and Rosemary Grant, and Nils Ryman for comments on a draft version. The National Genomics Infrastructure (NGI)/Uppsala Genome Center provided service in massive parallel sequencing, and the computational infrastructure was provided by the Swedish National Infrastructure for Computing (SNIC) at UPPMAX, partially funded by the Swedish Research Council through grant agreement no. 2018-05973.

## Author contributions

L.A. conceived the study. J.G., M.E.P. and A.W performed bioinformatic analysis. U.B., J.L. and L.W. collected herring samples and contributed to the interpretation of data. A.C. constructed sequencing libraries. B.D. performed morphological analysis. Y.W. performed otolith analysis. M.K. and H.W. were responsible for the analysis of organochlorine content. L.A., J.G., M.P. wrote the paper with input from all authors. All authors read and approved the final version.

## Supplementary Materials

**Extended Data Fig. 1.**
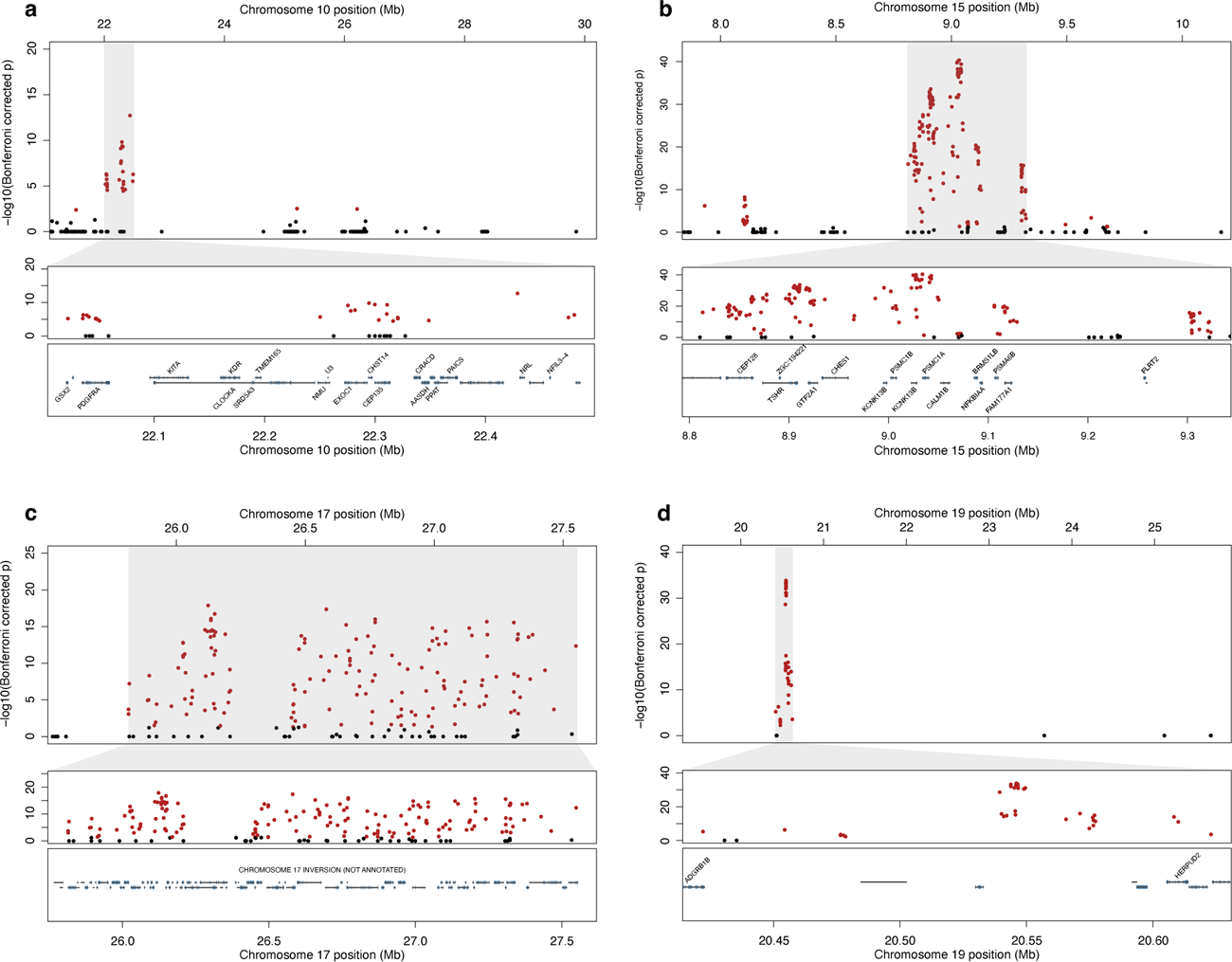
Summary of genome-wide, SNP-by-SNP, contrasts between spring-spawning Baltic herring and Slåttersill for the four main genomic regions on chromosomes 10 (**a**), 15 (**b**), 17 (**c**), and 19 (**d**). The topmost insets visualize chromosome-specific contrasts across the chromosome, while the center and bottommost insets visualize a zoomed version of the peak region of interest (denoted by a grey box). Gene annotations intersecting with regions of interest are annotated in the bottommost inset, while points highlighted in red denote SNP with significant *P*-values (<0.05) following Bonferroni correction. A list of all significant genes in the above plots is available from **Supplementary Table S3**. Chromosome 17 was not annotated as the region of interest spans the entire inversion region.

**Extended Data Fig. 2.**
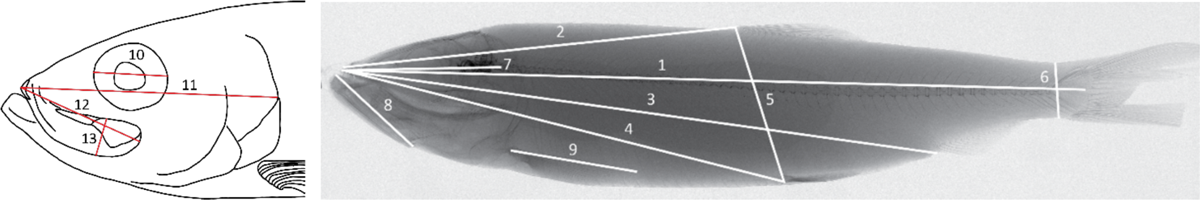
Measurements taken on *Clupea harengus* specimens; 1 standard length (SL), from upper jaw symphysis to middle base of caudal fin; 2, predorsal length from upper jaw symphysis to origin of dorsal fin; 3, preanal length, from upper jaw symphysis to origin of anal fin; 4, prepelvic length, from upper jaw symphysis to origin of pelvic fin; 5, body depth, origin of dorsal fin to origin of pelvic fins; 6, depth of caudal peduncle at origins of caudal fin procurrent fin rays; 7, cranium length, from upper jaw symphysis to articulation between the Atlas and next vertebra; 8, lower jaw length, from symphysis of dentary to retroarticular; 9, Pectoral fin length from base of first ray to tip of longest ray, digital calipers. 10, orbital horizontal diameter, x-ray; 11, head length, from upper jaw symphysis to posterior tip of operculum; 12, upper jaw length, from symphysis of premaxilla to posterior end of maxilla; 13, upper jaw depth, as greatest depth of maxilla and supramaxilla.

**Extended Data Fig. 3.**
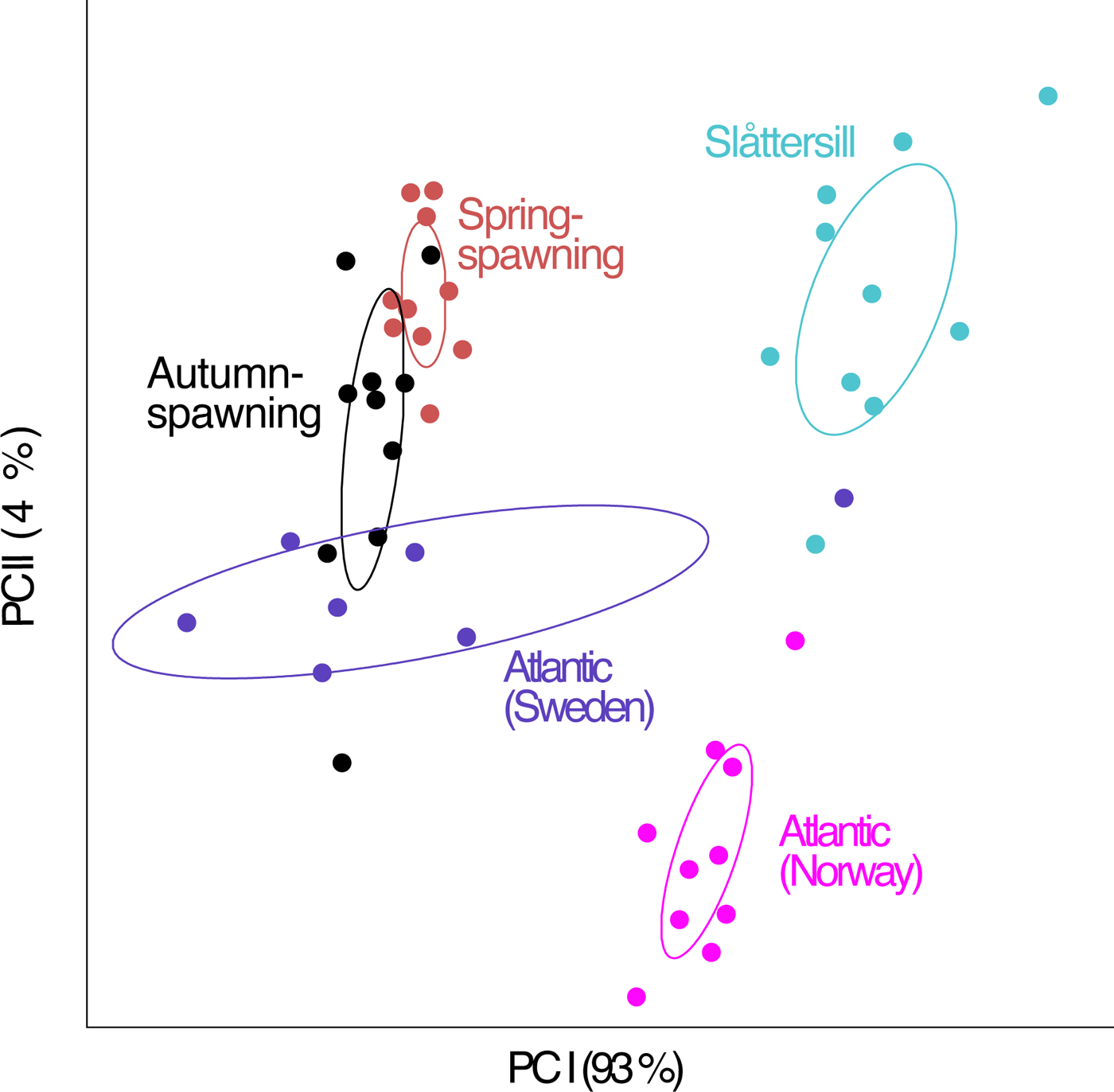
Principal component analysis of size and shape summarizing morphometric variation based on log-transformed measurement obtained from 47 specimens of *Clupea harengus* as shown in **Supplementary Table 4**. Corresponding character loadings are given in **Supplementary Table 5**. Ellipses indicate centroid positions with 95% confidence interval.

**Extended Data Fig. 4.**
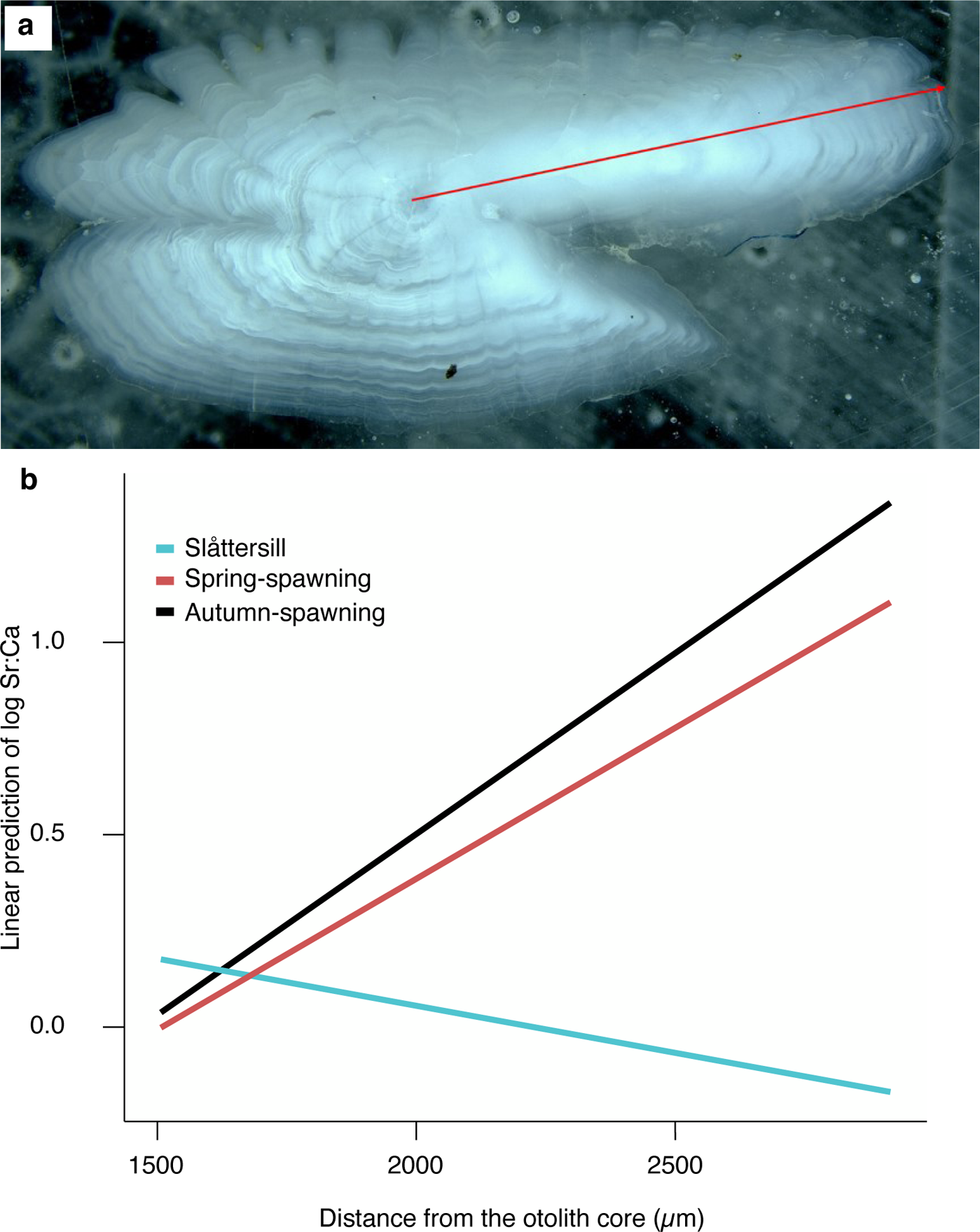
Results of otolith analysis. (**a**) Illustration of line transect (red line) from the otolith core to the anterior rostrum edge (top panel) used in lifelong otolith strontium:calcium profiling. (**b**) Pairwise comparison of otolith log transformed Sr:Ca slopes of Slåttersill (blue), autumn-spawning Baltic herring (black) and spring-spawning Baltic herring (red) from Gävlebukten. Only data from ≥ 1500 µm from the core were used in the analysis, corresponding to otolith material from adult herring only.

**Extended Data Fig. 5.**
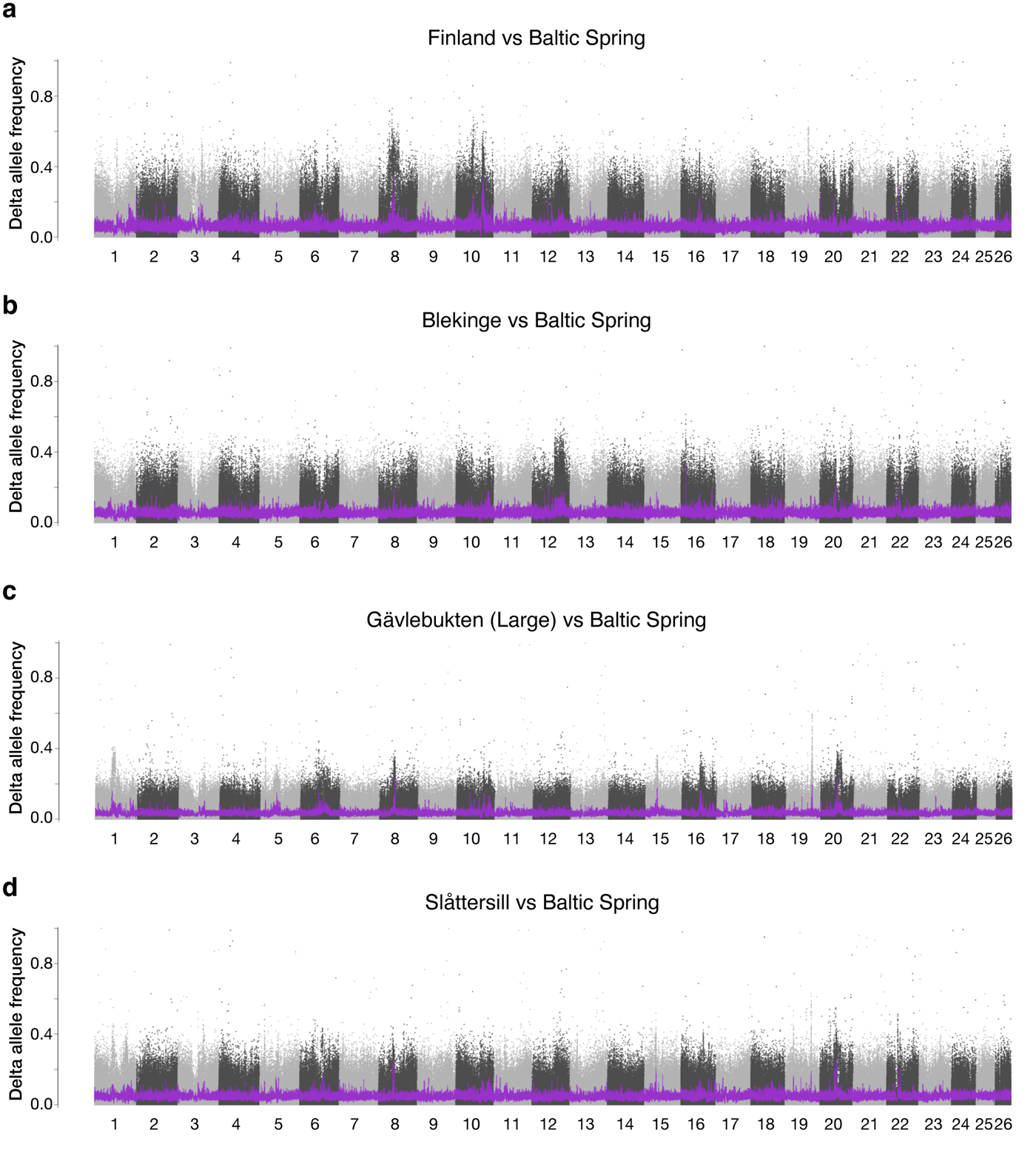
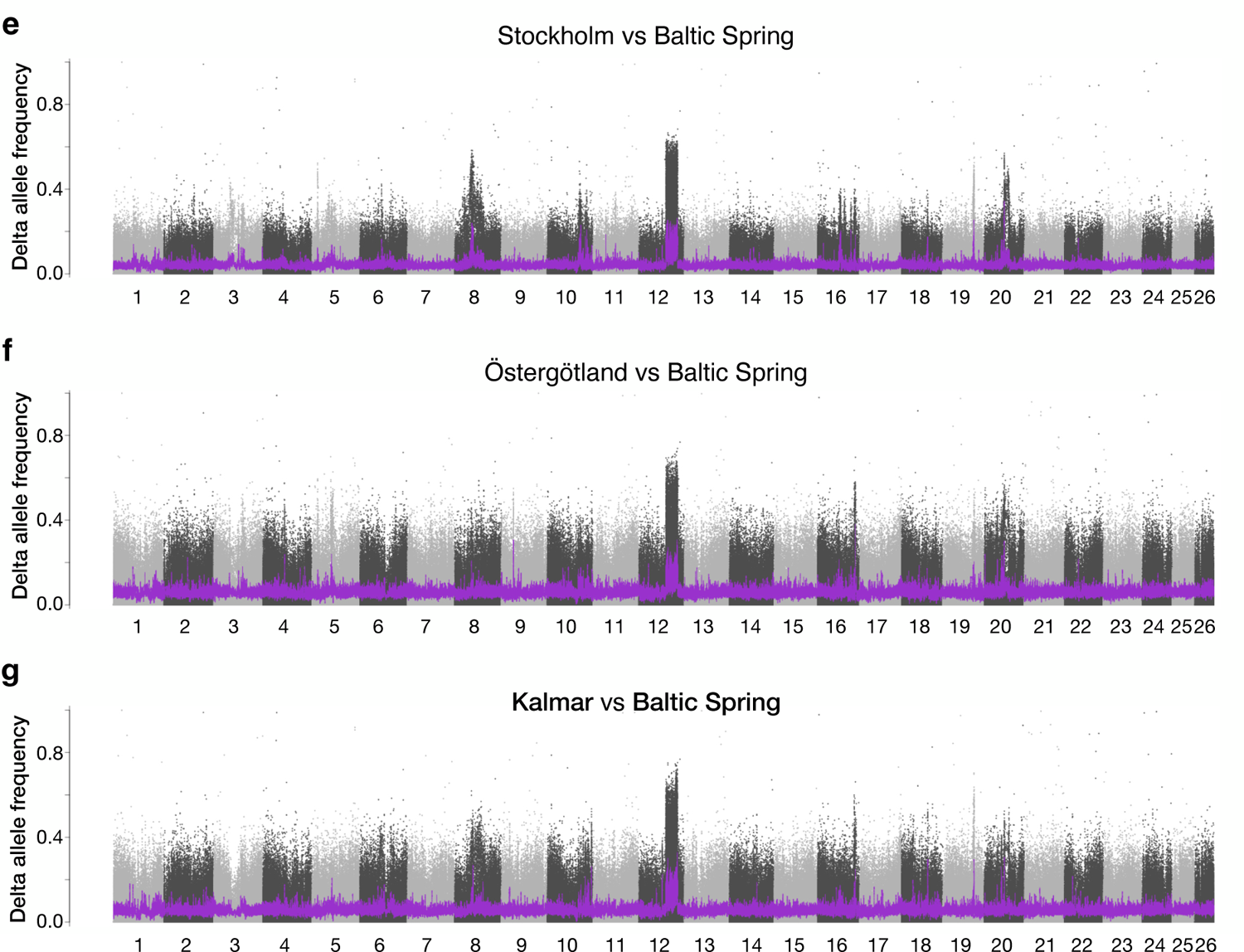
Pairwise delta allele frequency contrasts between small spring-spawning Baltic herring and Finland (**a**), Blekinge (**b**), Gävlebukten (Large, **c**), Slåttersill (**d**), Stockholm (**e**), Östergötland (**f**), and Kalmar (**g**). Notably, plots **a – d** represent large herring that exhibit a Northern haplotype at chr12, while plots **e – g** represent large herring with the Southern chr12 haplotype.

**Extended Data Fig. 6.**
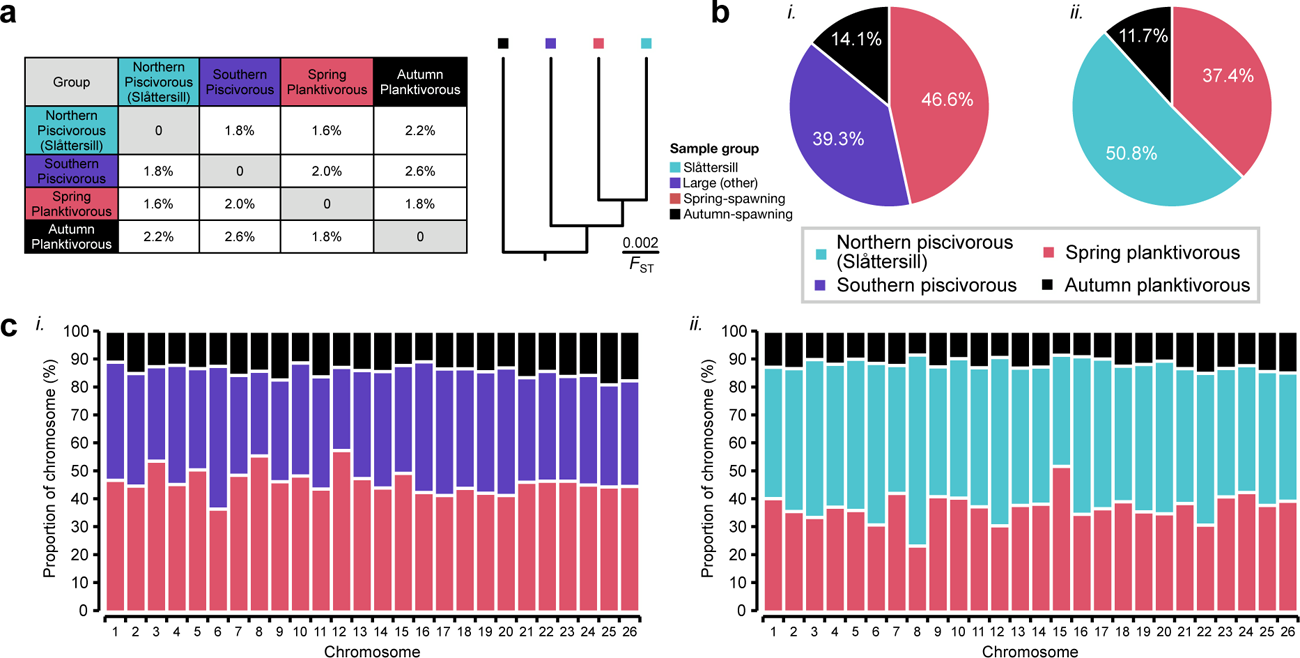
Pairwise genetic divergence among Baltic herring ecotypes. (**a**) Pairwise *F*_ST_-matrix (left) and corresponding UPGMA clustering (right). Comparing northern piscivorous herring (*i*) and southern piscivorous herring (*ii*), respectively, the pie charts (**b**) show the overall proportion of 10-kbp windows across chromosomes 1–26 (n=79,406) with the lowest *F*_ST_ between each piscivorous population and each of the other ecotypes, and the stacked bar plots (**c**) show the corresponding signature on a chromosome-by-chromosome basis.

**Supplementary Table 1.**
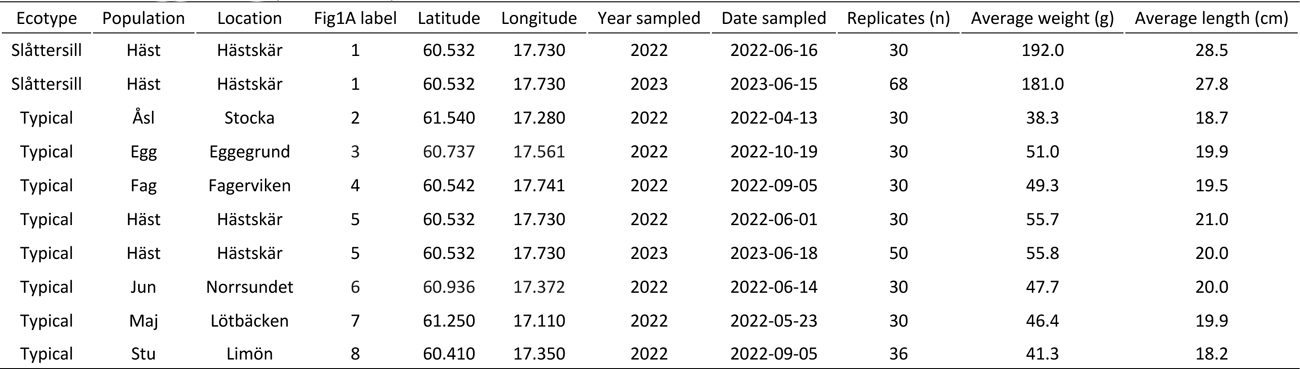
Summary of Baltic herring sampled from southern Bothnian Sea in 2022 and 2023 used for SNP genotyping on the MultiFishSNPChip_1.0 array (FSHSTK1D).

| Ecotype | Population | Location | Fig1A label | Latitude | Longitude | Year sampled | Date sampled | Replicates (n) | Average weight (g) | Average length (cm) |
| --- | --- | --- | --- | --- | --- | --- | --- | --- | --- | --- |
| Slåttersill | Häst | Hästkär | 1 | 60.532 | 17.730 | 2022 | 2022-06-16 | 30 | 192.0 | 28.5 |
| Slåttersill | Häst | Hästkär | 1 | 60.532 | 17.730 | 2023 | 2023-06-15 | 68 | 181.0 | 27.8 |
| Typical | Åsl | Stocka | 2 | 61.540 | 17.280 | 2022 | 2022-04-13 | 30 | 38.3 | 18.7 |
| Typical | Egg | Eggegrund | 3 | 60.737 | 17.561 | 2022 | 2022-10-19 | 30 | 51.0 | 19.9 |
| Typical | Fag | Fagerviken | 4 | 60.542 | 17.741 | 2022 | 2022-09-05 | 30 | 49.3 | 19.5 |
| Typical | Häst | Hästkär | 5 | 60.532 | 17.730 | 2022 | 2022-06-01 | 30 | 55.7 | 21.0 |
| Typical | Häst | Hästkär | 5 | 60.532 | 17.730 | 2023 | 2023-06-18 | 50 | 55.8 | 20.0 |
| Typical | Jun | Norrsundet | 6 | 60.936 | 17.372 | 2022 | 2022-06-14 | 30 | 47.7 | 20.0 |
| Typical | Maj | Lötbäcken | 7 | 61.250 | 17.110 | 2022 | 2022-05-23 | 30 | 46.4 | 19.9 |
| Typical | Stu | Limön | 8 | 60.410 | 17.350 | 2022 | 2022-09-05 | 36 | 41.3 | 18.2 |

**Supplementary Table 2.** Summary of age estimates based on the analysis of otoliths.

|  | *N* | Age at capture |  |
| --- | --- | --- | --- |
| | | Mean (± SD) | *P* * |
| Slåttersill | 10 | 7.2 (± 3.4) | - |
| Spring-spawning Baltic herring | 10 | 10.1 (± 3.2) | 0.001 |
| Autumn-spawning Baltic herring | 10 | 9.2 (± 2.3) | 0.006 |

**Supplementary Table 3**. Catalog of all significant loci (Bonferroni corrected *P* < 0.05) in a SNP-by-SNP contrast between Slåttersill and spring-spawning Baltic herring. (Excel file)

**Supplementary Table 4**. Samples of Atlantic and Baltic herring included in the morphological analyses. Summary of the analysis of meristic characters. (Excel file)

**Supplementary Table 5.** Character loadings on principal component I-V for 13 morphological measurements taken on 47 specimens of *Clupea harengus*.

|  | I | II | III | IV | V |
| --- | --- | --- | --- | --- | --- |
| Standard length | 0.183 | -0.014 | 0.004 | 0.002 | 0.013 |
| Orbital horizontal diameter | 0.087 | 0.055 | -0.017 | 0.000 | 0.009 |
| Head length | 0.150 | 0.020 | 0.005 | -0.018 | -0.008 |
| Cranium length | 0.136 | 0.053 | -0.012 | 0.012 | -0.001 |
| Body depth | 0.239 | -0.047 | -0.035 | 0.011 | -0.024 |
| Caudal peduncle depth | 0.216 | -0.060 | 0.013 | -0.005 | 0.014 |
| Predorsal length | 0.183 | -0.006 | -0.001 | -0.001 | 0.012 |
| Preanal length | 0.189 | -0.009 | 0.001 | 0.009 | 0.008 |
| Prepelvic length | 0.180 | 0.005 | 0.001 | 0.012 | 0.003 |
| Lower jaw length | 0.147 | 0.045 | 0.003 | 0.007 | 0.008 |
| Upper jaw length | 0.148 | 0.034 | -0.002 | -0.014 | -0.009 |
| Upper jaw height | 0.136 | 0.013 | 0.051 | 0.012 | -0.019 |
| Pectoral fin length | 0.174 | 0.003 | -0.001 | -0.030 | -0.004 |
| Variance explained by components | 0.380 | 0.016 | 0.004 | 0.002 | 0.002 |
| Percent of total variance explained | 93.3 | 3.9 | 1.1 | 0.5 | 0.4 |

**Supplementary Table 6.** Summary of biochemical comparison of Slåttersill (n=98) and spring-spawning Baltic herring (n=170) sampled from Gävlebukten across 2022 and 2023. All *P* values were derived from t-test comparing Slåttersill and spring-spawning Baltic herring.

| | Mean (± SD) | | |
| --- | --- | --- | --- |
|  | Slåttersill | Spring-spawning<br>Baltic herring | *P*-value |
| *Pollutant analysis* |  |  |  |
| Fat content (%) | 10.0 (± 2.1) | 3.9 (± 0.6) | 0.001 |
| ΣPCDD/F (pg TEQ/g ww) | 2.2 (± 0.5) | 5.2 (± 0.8) | <0.001 |
| ΣPCDD/F + dl-PCB (pg TEQ/g ww) | 5.2 (± 1.0) | 8.6 (± 1.7) | <0.001 |
| ΣPCB6 (ng/g ww) | 33.2 (± 7.3) | 40.4 (± 6.4) | 0.05 |
| *Isotope analysis* |  |  |  |
| N (%) | 9.9 (± 0.9) | 11.8 (± 1.0) | 0.003 |
| $\delta^{15}\text{N}$ (‰) | 10.5 (± 0.3) | 10.4 (± 0.4) | 0.34 |
| F<sub>N</sub> (%) | 0.4 (± 0.0) | 0.4 (± 0.0) | 0.34 |
| C (%) | 57.2 (± 1.7) | 53.2 (± 2.7) | 0.003 |
| $\delta^{13}\text{C}$ (‰) | -24.3 (± 0.6) | -22.8 (± 0.7) | 0.001 |
| F<sub>C</sub> (%) | 1.1 (± 0.0) | 1.1 (± 0.0) | 0.001 |
| C/N ratio | 5.8 (± 0.7) | 4.5 (± 0.6) | 0.004 |

**Supplementary Table 7.**
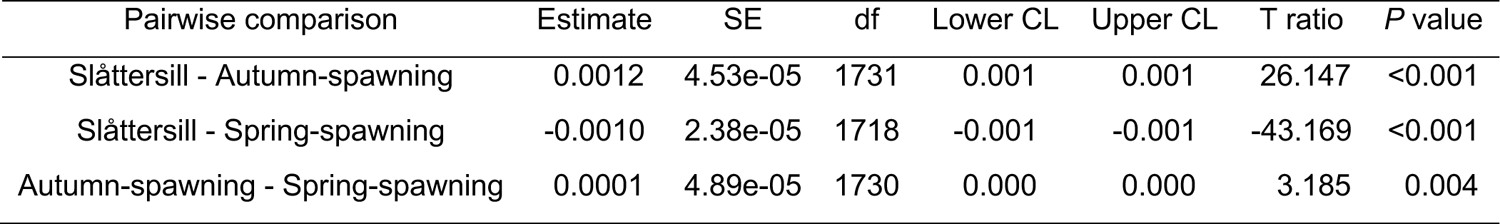
Results of pairwise comparison of otolith log transformed Sr:Ca slopes (≥ 1500 µm from the core) of Slåttersill, spring- and autumn spawning herring from Gävlebukten (Extended Data Fig. 4b). CL = confidence level.

**Supplementary Table 8.**
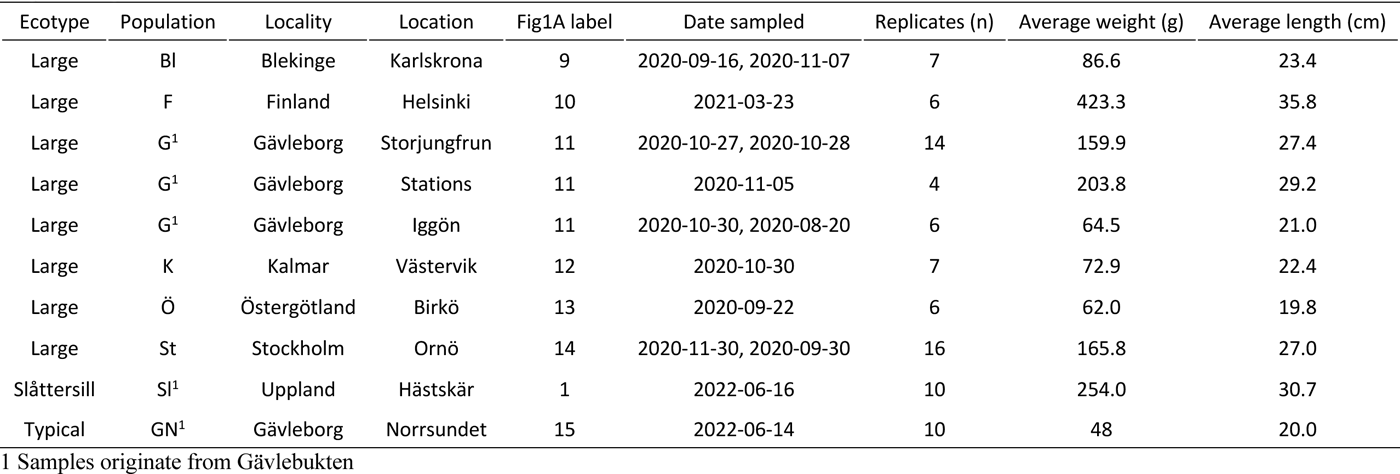
Summary of large herring and control sample (Gävleborg, Norrsundet) sampled from the Baltic Sea for whole genome sequencing.

**Supplementary Table 9.** Levels of genetic variation across the genome (chromosomes 1 to 26) and estimated effective population. The analysis is based on the assumption that all sites along chromosomes were accessible (n=724,810,791 bp). Long-term inference of effective population size (*N_e_*) estimated assuming mutation-drift equilibrium in an idealized population.

| Group | Sample size (n) | SNPs (n) | $\theta_w$/bp (%) | $\pi$/bp (%) | Tajima's D | $N_e$ (n) |
| --- | --- | --- | --- | --- | --- | --- |
| Spring planktivorous<sup>1</sup> | 80 | 16,944,883 | 0.47 | 0.21 | -1.92 | 590,007 |
| Autumn planktivorous<sup>2</sup> | 60 | 15,266,334 | 0.45 | 0.22 | -1.84 | 564,593 |
| Northern piscivorous<sup>3</sup> | 68 | 17,268,431 | 0.50 | 0.24 | -1.86 | 621,815 |
| Southern piscivorous<sup>4</sup> | 58 | 15,098,960 | 0.45 | 0.23 | -1.78 | 562,528 |

1 Grouping of samples:
Gvl-25 – 34; Sth02 – 3,6,12,22,30,38,41,43,46,50; Sth07 – 3:6,14:16,20,24,26; Upp05 – 1,2,11:14,16,19,23,25
2 Grouping of samples:
Gav18 – 3,12,17,23,27,30,39,40,46,47; Gav19 – 13,14,16,21:23,25,28:30; Gav20 – 3,7,24,50,53,63,70:74
3 Grouping of samples:
Gvl – 1:24; Upp – 1:10
4 Grouping of samples:
Klm – 1:7; Ogt – 1:6; Sth – 1:16

**Supplementary Table 10**. Full list of SNP designs for the MultiFishSNPChip_1.0 array (FSHSTK1D). The seventyonemer column illustrates the target SNP (enclosed within square bracket) for a given position, and its flanking regions which may contain additional known SNPs (enclosed in round brackets). The chromosomal positions refer to *Clupea harengus* reference assembly (Ch_v2.0.2.fasta). (Excel file)

